# The Upstream Sequence Transcription Complex Dictates Nucleosome Positioning and Promoter Accessibility at piRNA Genes in the *C. elegans* Germ Line

**DOI:** 10.1101/2023.05.10.540274

**Authors:** Nancy Sanchez, Lauren E Gonzalez, Valerie Reinke

## Abstract

The piRNA pathway is a conserved germline-specific small RNA pathway that ensures genomic integrity and continued fertility. In *C. elegans* and other nematodes, Type-I piRNA precursor transcripts are expressed from over 10,000 small, independently regulated genes clustered within two discrete domains of 1.5 and 3.5 MB on Chromosome IV. These large clusters likely play a significant role in promoting germline-specific expression of piRNAs, but the underlying mechanisms are unclear. By examining the chromatin environment specifically in isolated germ nuclei, we demonstrate that piRNA clusters are located in closed chromatin, and confirm the enrichment for the inactive histone modification H3K27me3. We further show that the piRNA biogenesis factor USTC (Upstream Sequence Transcription Complex) plays two roles – it promotes a strong association of nucleosomes throughout the piRNA clusters, and it organizes the local nucleosome environment to direct the exposure of individual piRNA genes. Overall, this work reveals new insight into how chromatin state coordinates transcriptional regulation over large genomic domains, which has implications for understanding global genome organization in the germ line.

## INTRODUCTION

PIWI-interacting RNAs (piRNAs) are part of a conserved small RNA pathway that maintains germline integrity and fidelity in many animal species. piRNAs silence aberrant expression of foreign genetic elements by associating with the PIWI sub-family of Argonaute proteins (LEE *et al*. 2012; SHEN *et al*. 2018) and targeting transcripts via antisense complementarity, which triggers co-transcriptional or post-transcriptional repression mechanisms (OZATA *et al*. 2019). In *C. elegans*, there are over 10,000 sequence-diverse piRNAs (RUBY *et al*. 2006; BATISTA *et al*. 2008). These piRNAs are 21 nucleotides long at maturity, have a strong 5′ uridine bias, and are uniquely expressed in the germline.

*C. elegans* piRNAs are encoded as individual transcription units and transcribed by a paused form of RNA polymerase II into ∼28 or ∼48 nucleotide precursors (RUBY *et al*. 2006; BATISTA *et al*. 2008; BELTRAN *et al*. 2021). The two types of *C. elegans* piRNAs are defined by genome organization: Type-I piRNAs are encoded within two large clusters on chromosome IV spanning 2.5 Mb and 3.7 Mb, whereas individual Type-II piRNAs are located throughout the genome, often near promoters of coding genes (RUBY *et al*. 2006; GU *et al*. 2012; BILLI *et al*. 2013). Clustering of piRNA genes is conserved across nematode species (SHI *et al*. 2013; BELTRAN *et al*. 2019), implying that clustering facilitates expression. The transcription of Type-I, but not Type-II, piRNA precursors requires the Upstream Sequence Transcription Complex (USTC) comprising PRDE-1, SNPC-4, TOFU-4, and TOFU-5 (KASPER *et al*. 2014; WEICK *et al*. 2014; WENG *et al*. 2019). Intriguingly, both SNPC-4 and TOFU-4 contain SANT protein domains, suggesting that they have the capacity to bind histones or histone modifications (BOYER *et al*. 2004; WENG *et al*. 2019). The USTC coats piRNA gene clusters broadly, with preferential binding at the piRNA-specific Ruby motif upstream of piRNA promoters (RUBY *et al*. 2006; WENG *et al*. 2019). Each component of the complex must be present for the others to bind within piRNA clusters (KASPER *et al*. 2014; WENG *et al*. 2019). While SNPC-4 and TOFU-5 have been implicated in other functions and have binding sites throughout the genome, TOFU-4 and PRDE-1 bind and function primarily at piRNA clusters, providing germline specificity (KASPER *et al*. 2014; WENG *et al*. 2019; HOU *et al*. 2022). The precise mechanisms by which USTC promotes piRNA expression are incompletely understood.

According to genomic data from whole worms, the chromosome IV piRNA clusters are enriched for the repressive H3K27me3 mark (BELTRAN *et al*. 2019). piRNA transcription requires both H3K27me3 methyltransferases and the H3K27me3 reader UAD-2 (USTC Association Dependent 2) (BELTRAN *et al*. 2019; HUANG *et al*. 2021), suggesting that a repressive chromatin environment is important for piRNA transcription. However, the chromatin environment of piRNA clusters specifically within germline cells, and how that environment facilitates piRNA transcription, are still unclear.

Here, we use genomic analyses of isolated germ nuclei (IGN) to specifically interrogate the chromatin environment of *C. elegans* piRNA clusters in germ cells, and uncover a role for USTC in affecting chromatin organization to drive Type-I piRNA transcription. We show that the piRNA clusters are indeed enriched for H3K27me3 in the germline, and reveal that chromatin accessibility is locally modulated around piRNA transcription start sites (TSSs) by USTC. By probing chromatin accessibility as well as nucleosome and histone H3 positioning, we found a well-defined nucleosome just upstream of the piRNA TSS whose positioning and stability depends on USTC. Together, our data show that in germ cells, piRNA clusters have an organized nucleosome landscape with high nucleosome density and high levels of the H3K27me3 silencing modification. Within this repressive domain, UTSC establishes organized nucleosomes that permit local transcription of individual piRNA genes. This structured micro-environment within a larger repressive domain likely facilitates RNA pol II pausing, limiting transcription elongation and encouraging production of short piRNA precursor transcripts. This mechanism therefore coordinates local and long-range regulation of thousands of noncoding RNA genes in a tissue-specific manner.

## RESULTS

### piRNA gene promoters specifically lack H3K27me3 within H3K27me3-enriched domains

Previous studies examining chromatin profiles from whole animal datasets found that the piRNA gene clusters are enriched for the repressive histone modification H3K27me3 (BELTRAN *et al*. 2019). Because prior analyses included both somatic tissues as well as the germline, we wished to determine the overall chromatin status of the piRNA clusters specifically in germ cells. We therefore isolated germline nuclei (IGN) from wild type adults and performed chromatin immunoprecipitation followed by sequencing (ChIP-seq) for three histone modifications that have known significant effects on germline gene expression: the active histone modification H3K36me3, and the repressive histone modifications H3K9me3 and H3K27me3. Consistent with the previously described whole animal datasets, the piRNA cluster is enriched for H3K27me3 in IGN (Figure 1A). Strikingly, within this enriched domain, a strong local depletion of H3K27me3 was apparent at individual piRNA genes; this pattern was much more detectable in IGN than in whole animals, suggesting that it is specific to the germ line (Figure 1B,C). By contrast, both H3K36me3 and H3K9me3 levels do not change significantly across piRNA domains in IGN of either wild type or *prde-1* mutants compared to whole animals (Supplemental Figure 1A-C). A local depletion of H3K9me3 at piRNA genes similar to that seen for H3K27me3 was also detectable, although the pattern was much weaker (Supplemental Figure 1B). These data indicate that the IGN technique provides higher resolution to capture germline-specific chromatin profiles, and shows that while the broader piRNA clusters are generally enriched for H3K27me3, the immediate neighborhood surrounding individual piRNA genes is not.

**Figure 1.**
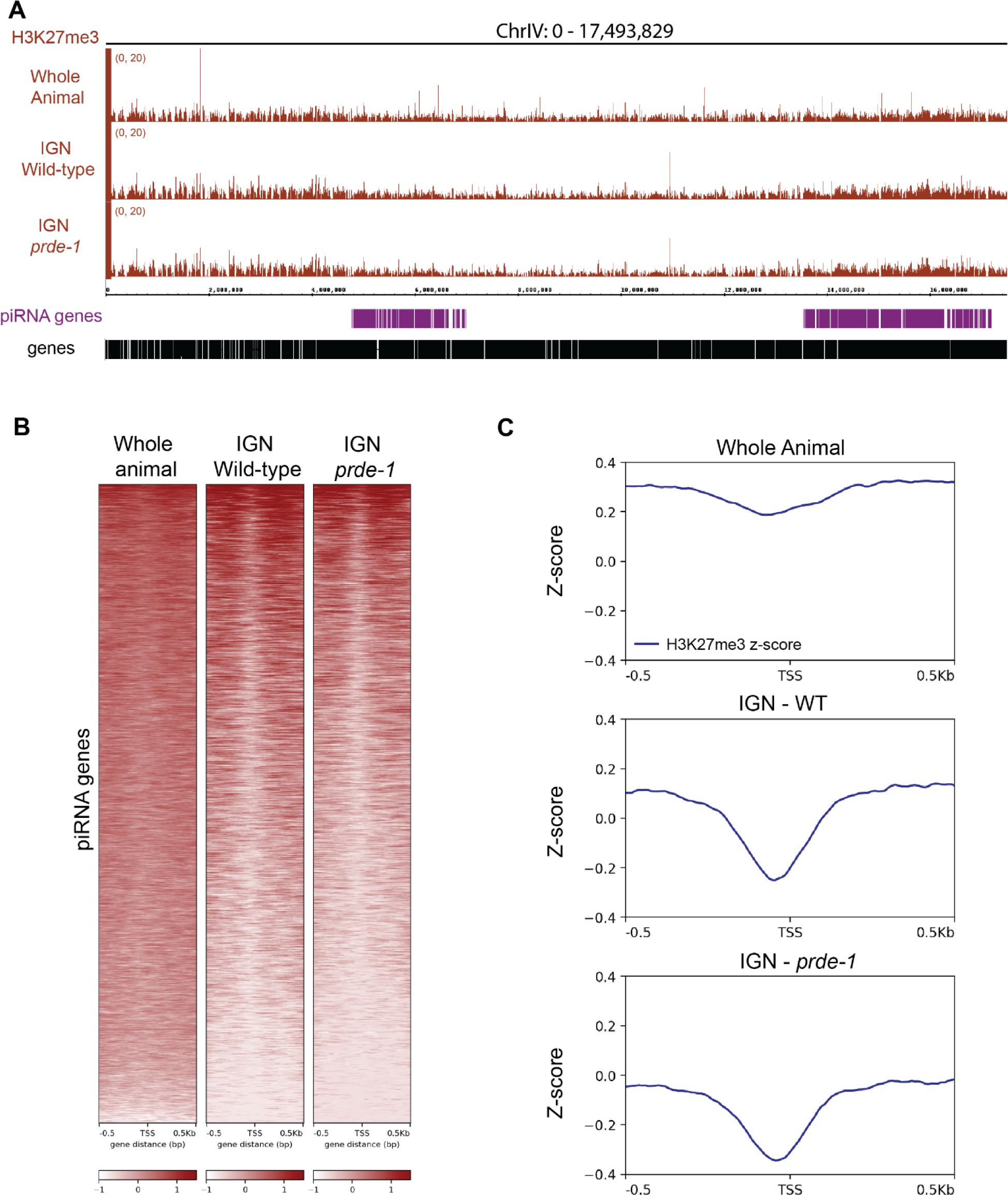
piRNA genes are depleted for H3K27me3 in germ nuclei. A) Genome browser view of H3K27me3 patterns across chromosome IV for whole animal, wild type IGN, and *prde-1* mutant IGN. B) Heatmap of H3K27me3 levels centered on the transcription start site (TSS) of piRNA genes in piRNA clusters. The H3K27me3 signal is represented as Z-scores. C) Metagene profile of H3K27me3 Z-score values across piRNA genes (1kb, centered on the piRNA TSS).

To investigate whether USTC plays a role in establishing and/or maintaining the histone modification landscape of the piRNA loci, we performed IGN-ChIP-seq for H3K36me3, H3K27me3, and H3K9me3 in the *prde-1* mutant background. PRDE-1 is a germline-specific component of USTC that is required for USTC assembly at piRNA clusters and production of piRNA precursor transcripts (KASPER *et al*. 2014; WEICK *et al*. 2014; WENG *et al*. 2019). Overall levels of H3K27me3 across the piRNA clusters are somewhat lower in the *prde-1* mutant relative to wild type (Figure 1B), but the relative depletion of H3K27me3 at individual piRNA genes persists (Figure 1C). We did not detect any significant genome-wide changes of H3K36me3 or H3K27me3 signal levels between wild type and the *prde-1* mutant genome wide (Supplemental Figure 2), although levels of H3K9me3 are very slightly increased in the *prde-1* mutant within piRNA clusters. In sum, these data show that USTC does not have a major influence the local patterns of histone modifications at individual piRNA genes, but does promote an overall higher level of H3K27me3 across the broader piRNA clusters.

### piRNA genes are located in low accessibility domains

piRNAs are abundantly expressed in the germ line, despite the enrichment of the repressive mark H3K27me3 throughout the piRNA clusters. However, the local depletion of H3K27me3 at individual piRNA genes (Figure 1B-C) suggests that this repressive environment might be disrupted locally to promote piRNA expression. We therefore investigated overall chromatin accessibility within the piRNA gene clusters using ATAC-seq (Assay for Transposase-Accessible Chromatin using sequencing (GRANDI *et al*. 2022)) in wild type and *prde-1* mutant IGN. Notably, chromatin accessibility was low overall throughout the piRNA clusters relative to the rest of the genome, and even lower in *prde-1* mutants (Figure 2A, inset). In wild type, many individual sites of chromatin accessibility that overlap with the piRNA clusters appear reduced in the *prde-1* mutant (Figure 2A). Genome-wide differential peak analysis demonstrates that the vast majority of accessible chromatin peaks were lost and very few gained in the *prde-1* mutant (Figure 2B). Specifically within piRNA clusters, differential peaks were only lost and none gained, indicating that the piRNA clusters become less accessible in the *prde-1* mutant. Moreover, most genes associated with the decreased ATAC peaks were protein coding genes; piRNA genes represented about 4% of the total (Supplemental Figure 3). Thus, the peaks lost in *prde-1* mutants are generally not associated with piRNA genes. Together these observations indicate that major changes in accessibility throughout the piRNA clusters are largely a consequence of changes to non-piRNA gene features in the region. This conclusion is supported by the fact that most of the open chromatin peaks lost in *prde-1* mutants are not restricted to the piRNA region but occur throughout the genome (Figure 2B).

**Figure 2.**
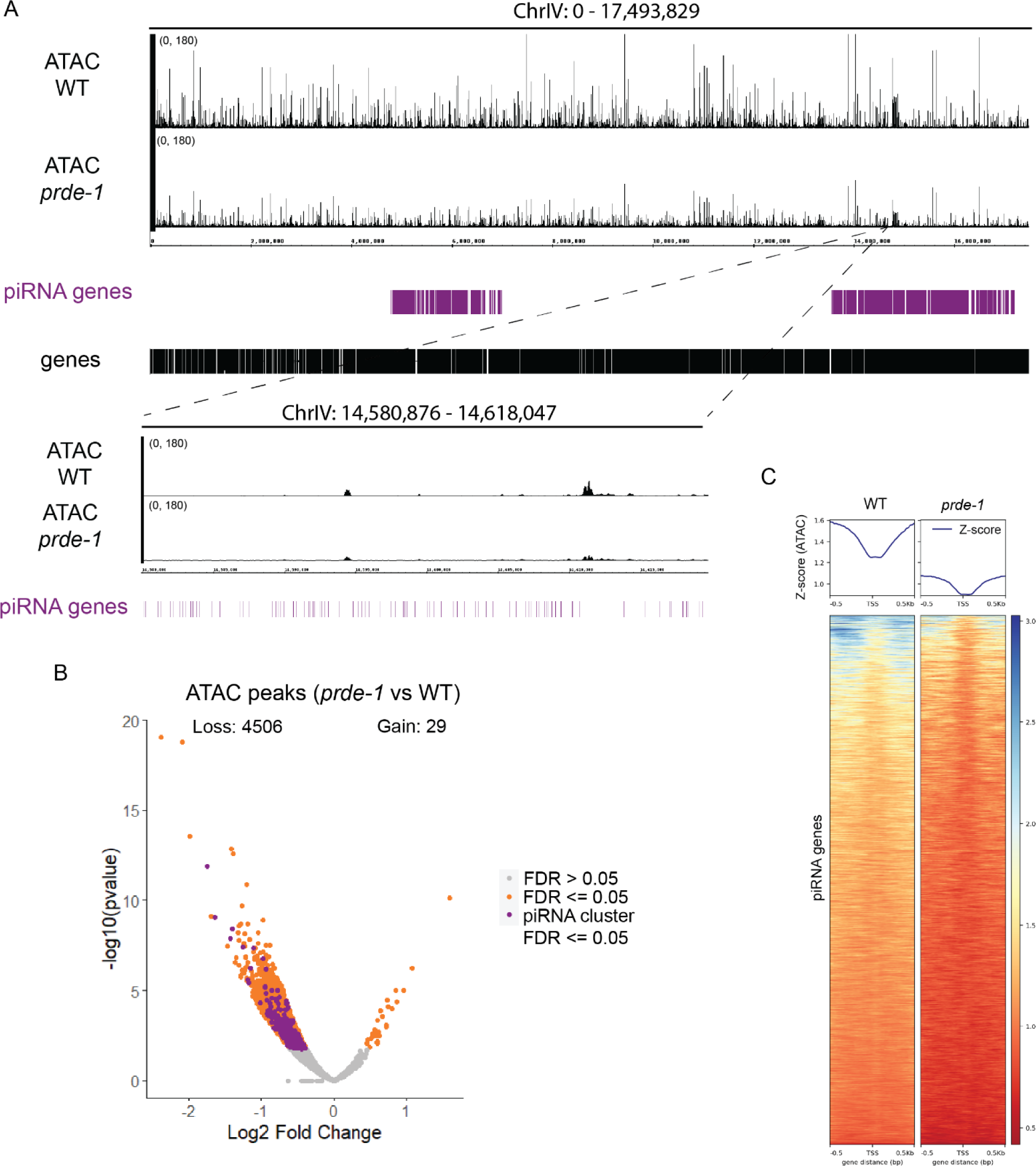
Chromatin accessibility is reduced at piRNA clusters in *prde-1* mutant germ nuclei. A) Genome browser view of ATAC signal in wild-type and *prde-1* mutant IGN across chromosome IV, with inset to show low signal in piRNA genes. B) Volcano plot representing differential chromatin accessibility peaks between *prde-1* mutant and wild type. Orange = statistically significantly different peaks (FDR <= 0.05). Grey = nonsignificant peaks (FDR > 0.05). Purple = differential peaks located in the piRNA gene cluster (including both piRNA genes and coding genes) that are statistically significant (FDR <= 0.05). C) Metagene plots and heatmaps displaying chromatin accessibility Z-score values in wild type and *prde-1* mutants (1kb, centered on the TSS of piRNA genes).

We next examined whether USTC affects local chromatin accessibility patterns around piRNA genes. In wild type, chromatin accessibility was low overall throughout the piRNA clusters relative to the rest of the genome, and even lower in *prde-1* mutants (Figure 2A, inset). This pattern was recapitulated at individual piRNA genes (Figure 2C). A subset of piRNA genes are located in relatively open chromatin regions (top few piRNAs in heatmap in Figure 2C), which leads to piRNA genes on average appearing to be less accessible than their surrounding sequences in the metagene plot(Figure 2C). Close inspection suggests an extremely subtle increase of accessibility at the piRNA transcription start site (TSS) relative to immediately adjacent sequences, which was diminished in the *prde-1* mutant (Figure 2D). Together, these results demonstrate that the piRNA clusters have relatively low overall accessibility, and the loss of USTC decreases regions of open chromatin genome-wide as well as specifically within the piRNA clusters.

### Local nucleosome organization at piRNA genes is dependent on USTC

High levels of H3K27me3 (Figure 1) and low accessibility (Figure 2) in the piRNA clusters of wild type IGN suggest a very dense nucleosome environment, so we next investigated local nucleosome organization at piRNA genes. To predict nucleosome occupancy and positioning, we used nucleoATAC, which selectively analyzes mononucleosome-sized fragments from ATAC-seq datasets (SCHEP *et al*. 2015). This analysis predicts that in wild type germ cells, well-defined nucleosomes are located immediately upstream and downstream of piRNA genes, with a depletion of a nucleosome precisely at the TSS (Figure 3A). To determine whether this pattern is affected by piRNA expression, we stratified piRNAs into deciles according to average transcript abundance from three publicly available datasets (GU *et al*. 2012; KASPER *et al*. 2014; BELTRAN *et al*. 2021). In wild type germ cells, predicted nucleosome patterns were well-defined for highly expressed piRNAs in the top decile, while those in the bottom decile had poorly-defined nucleosome patterns (Figure 3A). These observations are consistent with the depletion of H3K27me3 (Figure 1A-B), as well as the subtle increase in accessibility (Figure 2D) near the TSS of piRNA genes.

**Figure 3.**
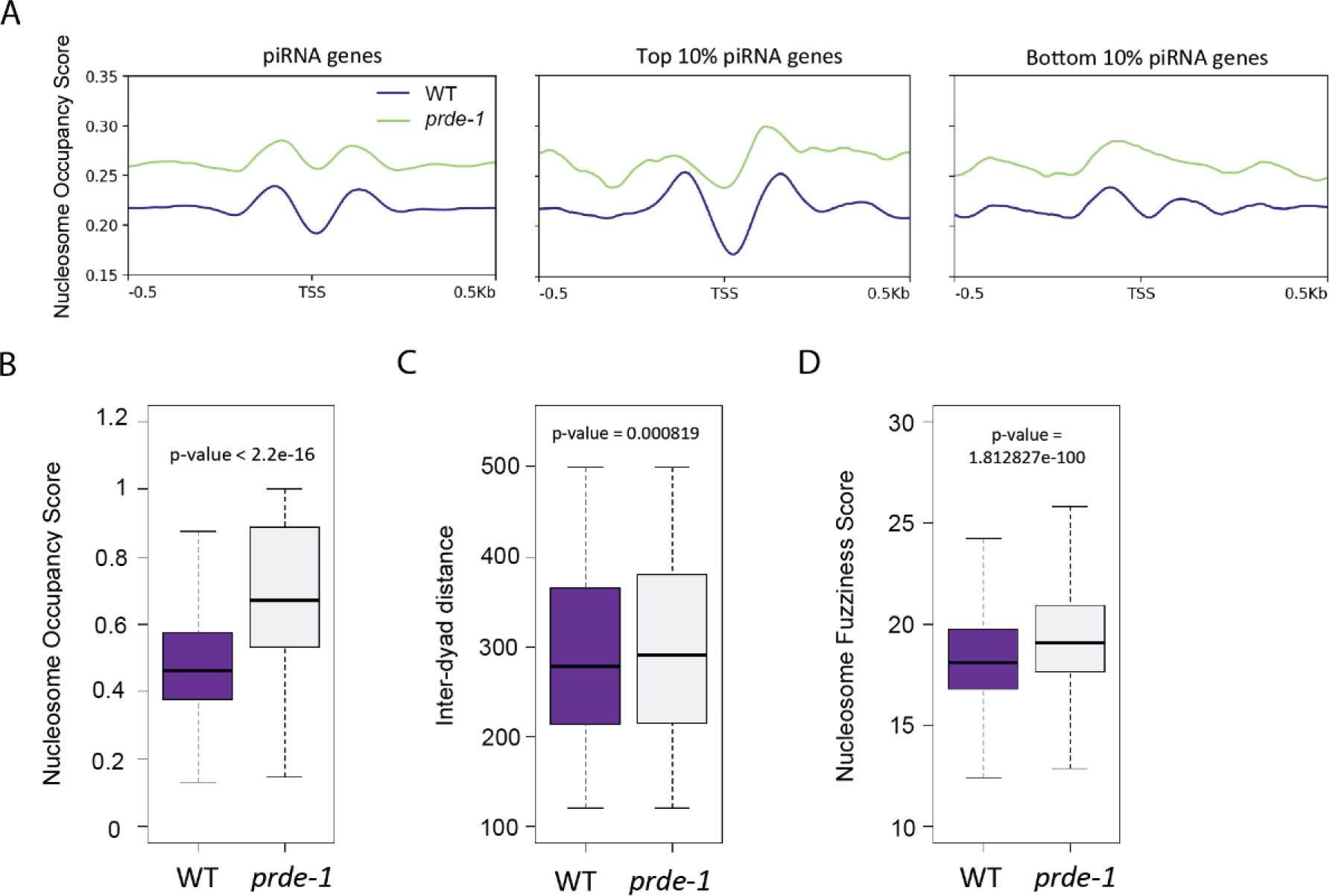
PRDE-1 contributes to nucleosome environment of piRNA genes. A) Metagene analysis of nucleosome occupancy scores (1kb, centered on the TSS of piRNA genes). Wild type values are in blue while *prde-1* mutant values are in green. Middle and right plot represent nucleosome occupancy scores for the top 10% expressed piRNA genes and bottom 10% expressed piRNA genes, respectively. B-D) Quantitative analysis of nucleosome occupancy scores, inter-dyad distance, and nucleosome fuzziness scores extracted from +/− 500 bp regions surrounding piRNA genes (measured from the TSS). Purple represents wild type and grey represents *prde-1* mutant. Statistical significance was calculated with Wilcoxon test.

In the *prde-1* mutant, we observed increased nucleosome occupancy near the promoters of piRNA genes compared to wild type (Figure 3A), which corresponds to the loss of chromatin accessibility at piRNA genes (Figure 2C). Regardless of piRNA abundance, overall occupancy is higher in the *prde-1* mutant, but the individual nucleosomes flanking the piRNA TSS are not as well-positioned as in wild type (Figure 3A). Notably, for highly expressed piRNA genes, the predicted upstream nucleosome in particular becomes much less well-defined in the *prde-1* mutant (Figure 3A, middle panel). Importantly, *prde-1*-dependent changes in predicted nucleosome patterns are not seen at genes with similar features of genomic organization and transcriptional regulation, such as genes with paused or docked pol II (MAXWELL *et al*. 2014) (Supplemental Figure 4). Together, these data suggest that distinct nucleosome environments may drive different levels of piRNA expression, and that USTC is necessary to establish a specific local organization of nucleosomes that flank the piRNA TSS.

By performing quantitative analysis of the nucleosome environment in 1kb regions around piRNA genes, we confirmed a significant increase of nucleosome occupancy in the *prde-1* mutant (Figure 3B). Additional measures of nucleosome organization reinforced these observations. Both inter-dyad distance, which is the distance between the centers of two neighboring nucleosomes, and nucleosome fuzziness score, which represents uncertainty in nucleosome positioning, were increased in *prde-1* mutants, suggesting that nucleosome placement is less well-defined when the USTC is disrupted (Figure 3C, D). Taken together, the NucleoATAC analysis supports a role for USTC in organizing nucleosome patterns in local regions around piRNA genes to establish an open configuration immediately adjacent to individual piRNA loci.

### USTC stabilizes histone H3 density across piRNA clusters

To independently assess the nucleosome environment at piRNA genes, we performed H3 ChIP-seq in wild type and *prde-1* mutant germline nuclei. Since the ATAC-seq was performed on live (uncrosslinked) IGN, we adapted Native ChIP-seq (BRIND’AMOUR *et al*. 2015) to assess H3 location under the same conditions (see Methods). Under standard salt conditions (150mM), we observed H3 enrichment throughout the piRNA gene clusters relative to the rest of the genome in both wild type and *prde-1* mutants (Figure 4A). On closer inspection, the H3 profile is anti-correlated with chromatin accessibility peaks, as expected (Figure 4B). Interestingly, a strong H3 signal apparent just upstream of the TSS at piRNA genes in wild type is diminished in the *prde-1* mutant (Figure 4C), suggesting that USTC maintains the upstream nucleosome at piRNA genes, consistent with the NucleoATAC analysis (Figure 3A).

**Figure 4.**
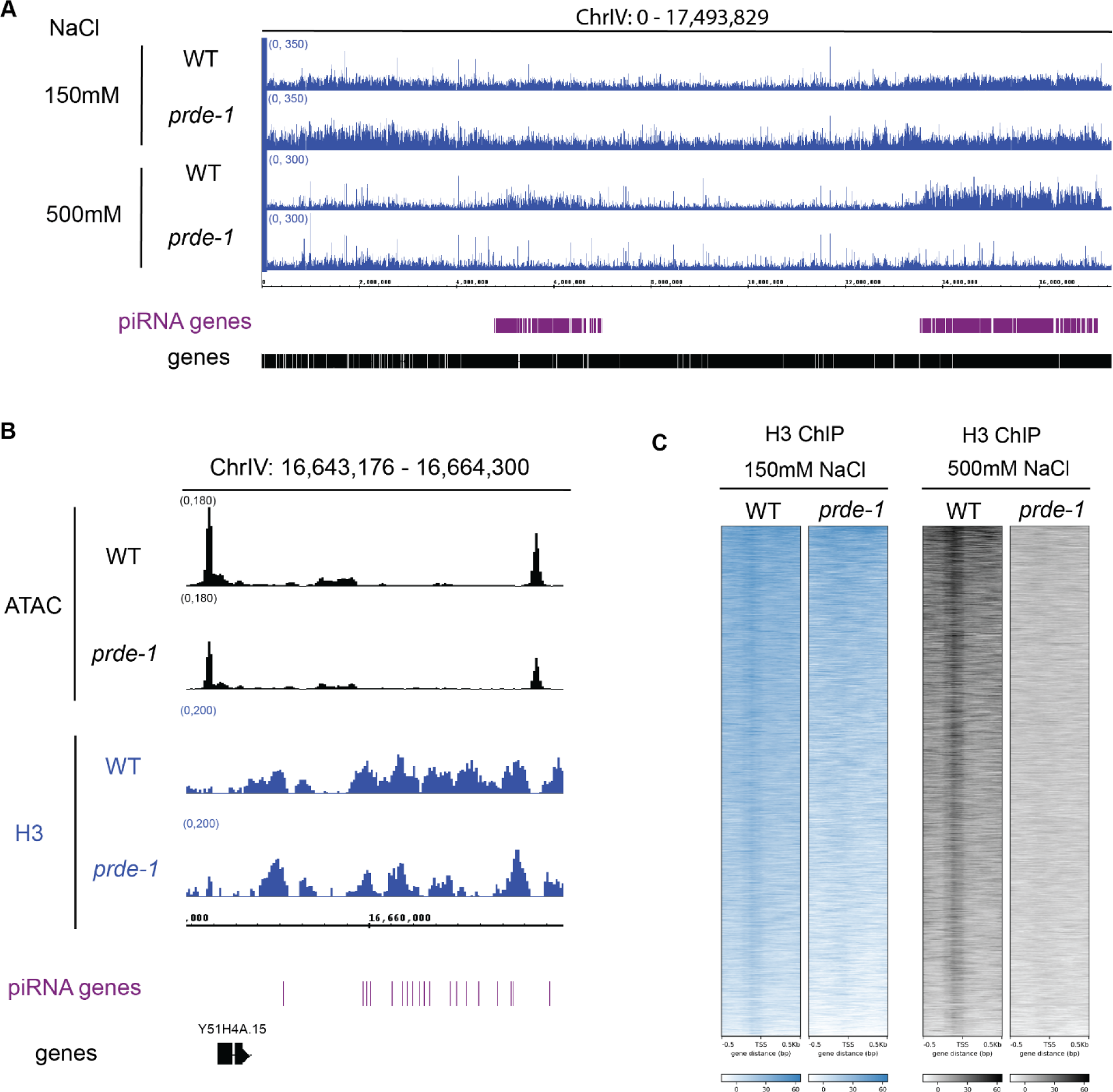
PRDE-1 maintains H3 enrichment across piRNA gene clusters. A) Genome browser view of Native H3 ChIP-seq signal (input subtracted) at low and high salt conditions. B) Genome browser view example of inverse relationship between wild type ATAC signal and wild type H3 ChIP signal (150mM salt condition). C) Heatmap representing wild-type and *prde-1* H3 signal of low salt (left) and high salt (right) conditions, 1kb, centered at the TSS of piRNA genes.

Because histone association with DNA is highly sensitive to salt concentration (SANDERS 1978; LI *et al*. 1993; JIN and FELSENFELD 2007; HENIKOFF *et al*. 2009; OOI *et al*. 2010), we assessed whether higher salt in the washes altered H3 profiles, as a measure of relative nucleosome affinity for DNA. In wild type IGN, increased salt drastically reduced H3 association with DNA throughout the genome, with the specific exception of the two piRNA domains, which retained strikingly high levels of H3 (Figure 4A). In the *prde-1* mutant, H3 levels were depleted across the piRNA gene clusters similarly to the rest of the genome, including complete loss of the strong H3 signal upstream of the piRNA gene TSS (Figure 4C). This observation indicates that USTC promotes strong association of nucleosomes with the DNA specifically at piRNA clusters. Together, analysis of H3 levels in IGN independently confirm the conclusions from the ATAC-seq data and nucleoATAC analysis: USTC both promotes the high overall nucleosome density across the piRNA gene cluster and establishes a structured nucleosome environment locally upstream of individual piRNA genes.

## DISCUSSION

In this report, we assess the specialized chromatin environment of the two genomic clusters of Type-I piRNA genes on Chromosome IV specifically in *C. elegans* germline nuclei, and determine the role of USTC in establishing this environment. We demonstrate that in the absence of USTC, the association of nucleosomes with DNA is less consistent and nucleosomes appear more mobile across piRNA clusters. Moreover, the loss of USTC function disrupts the predicted nucleosome positioning flanking the transcription start sites (TSSs) of individual piRNA genes, and the strength of this effect correlates with piRNA abundance. Thus, our data are consistent with a model in which USTC is necessary for the establishment of an organized chromatin environment in the germ line that facilitates piRNA expression.

Although it is difficult to determine precise nucleosome placement using genomic assays that rely on aggregate measurements, data from several independent experiments supports the conclusion that the piRNA TSS of is largely devoid of a nucleosome, but flanked by two well-positioned nucleosomes upstream and downstream. First, the H3K27me3 ChIP-seq data point to a local depletion of this modification immediately upstream and at the TSS (Figure 1C), which can be interpreted as either an unmodified or absent nucleosome. Second, nucleoATAC analysis of the ATAC-seq data predicts the absence of a nucleosome immediately at the TSS, with two well-positioned nucleosomes upstream and downstream (Figure 3A); the precision of this pattern correlates with abundance of piRNAs, suggesting that the pattern is related to piRNA transcription. Notably, the absence of a single nucleosome would expose approximately 147nt of DNA (not including linker sequence) (OOI *et al*. 2010), which is more than sufficient to encode the closely-spaced upstream regulatory motifs, promoter, and downstream motifs flanking each piRNA gene. Finally, Native ChIP-seq of H3, which does not distinguish between modified and unmodified H3, identifies an enrichment of H3 specifically upstream of the TSS. Importantly, in all three of these assays, loss of USTC function disrupts these patterns, suggesting that they are related to each other, as well as dependent on USTC.

Our data confirm that H3K27me3 is enriched throughout piRNA clusters in the germline, and reveal a strong local depletion of H3K27me3 at the TSSs of individual piRNA genes (Figure 1). This local H3K27me3 depletion was barely visible from whole animal data, indicating that genomic analyses in IGN greatly increase resolution and specificity. However, USTC does not seem to be directly involved in establishing these H3K27me3 patterns. Disrupting USTC function has no effect on the local depletion of H3K27me3 at piRNA TSSs and leads to only a mild reduction in H3K27me3 level across the piRNA clusters, possibly as a secondary consequence of the decreased stability of nucleosomes. Thus, the deposition of H3K27me3 is likely regulated by an independent mechanism. Indeed, the *C. elegans* orthologs of PRC2, a histone methyltransferase complex that methylates H3K27, as well as UAD-2, an H3K27me3 reader, are necessary for USTC binding at piRNA clusters (HUANG *et al*. 2021). Together, these observations suggest that deposition of H3K27me3 by PRC2 occurs independently of, and prior to, USTC binding. The use of heterochromatin to mark piRNA clusters and recruit specific regulatory factors for piRNA gene expression is not unique to *C. elegans*. In Drosophila, piRNA precursors located in dual-strand clusters are dependent on high levels of H3K9me3 at the promoter, which leads to the recruitment of the Rhino protein and subsequent transcription of the precursor RNA (RANGAN *et al*. 2011; LE THOMAS *et al*. 2014; MOHN *et al*. 2014; ANDERSEN *et al*. 2017). Thus, H3K27me3 and USTC in *C. elegans* may be analogous to H3K9me3 and Rhino in Drosophila. This similarity between species occurs despite profound differences in the two systems: Drosophila piRNAs largely encode repetitive, transposon-rich sequences, and are produced as large precursor RNAs that are subsequently cleaved into many piRNAs, while in *C. elegans*, piRNAs are incredibly sequence diverse and are independently transcribed from tiny individual genes (OZATA *et al*. 2019). The mechanistic reasons why both systems use heterochromatin-like states at piRNA genomic loci is unclear.

Overall, our data are consistent with a model in which USTC organizes a defined nucleosome environment around piRNA genes, which permits expression from a nucleosome-rich domain decorated with the repressive mark H3K27me3. Despite this effect on nucleosome positioning, no USTC protein component encodes an enzymatic domain that might actively remodel nucleosomes. Therefore, at this point, we cannot rule out that the nucleosome positioning seen at individual piRNA genes is a consequence of active RNA polymerase II, and that the role of USTC is to recruit RNA polymerase II, which then initiates transcription, leading to nucleosome displacement at the promoter and stabilization of upstream and downstream nucleosomes as a consequence. Alternatively, another nucleosome remodeler might be recruited by USTC to directly perform this function. Indeed, the ATPase-containing subunit of the Nucleosome Remodeling Factor (NURF) complex, ISW-1, is necessary for recruitment of USTC to piRNA clusters and for piRNA expression (HUANG *et al*. 2021). Future studies that incorporate candidate chromatin remodelers, as well as the RNA pol II machinery, will be required to better understand the precise mechanisms by which USTC coordinates piRNA expression in the germ line of *C. elegans*.

## MATERIALS AND METHODS

### C. elegans strains

Strains were maintained at 20°C on NGM plates seeded with OP50 (BRENNER 1974). VC2010 was used as a wild type strain, and the *prde-1(mj207)* strain was used as the *prde-1* null mutant strain (WEICK *et al*. 2014).

### Isolation of germline nuclei

Isolated germline nuclei (IGN) were harvested as in (HAN *et al*. 2019) with minor modifications. Worms were harvested at 49-52 hours for wild type and 56-59 hours for *prde-1(-),* when most animals had 4-6 embryos. Approximately 1.2 million worms were used per isolation. Worms were washed with 1x M9 in sets of 6 plates into a 50mL falcon tube, washed twice with M9, floated on sucrose, and washed again 3x in M9.

#### IGN for ChIP-seq

At the adult stage, worms were crosslinked for 30min in 2% formaldehyde diluted in M9. Formaldehyde was quenched by a 1 M Tris (pH 7.5) wash and washed twice with M9. Then the worms were washed with 10mL chilled Nuclei Purification Buffer (NPB) (50 mM HEPES pH 7.5, 40 mM NaCl, 90 mM KCl, 2 mM EDTA, 0.5 mM EGTA, 0.1% Tween 20, 0.2 mM DTT, 0.5 mM PMSF, 0.5 mM spermidine, 0.25 mM spermine, and 1x Complete Protease Inhibitor Cocktail). Worms were resuspended in 6mL of NPB and transferred into prechilled 7mL glass Dounce homogenizers (Wheaton cat. no. 06435A). Fifteen loose strokes were followed by 23 tight strokes with a quarter turn between each Dounce. The worms were transferred into chilled 50 mL Falcon tubes, and NPB was added to 10 mL. The tubes were vortexed on medium-high speed for 30 s, followed by 5 min on ice. The vortex and ice incubation were repeated. Worm debris was filtered with four 30μm strainers (MACS cat. no. 130-098-458) and seven 20 μm strainers (Pluriselect cat. no. 43-50020-03). Nuclei were spun at 3100 rpm for 6 min at 4°C. The supernatant was removed, and the nuclei resuspended in 1 mL NPB and transferred to a nonstick 1.5 mL tube (Ambion). A 5 μl aliquot was removed, incubated with DAPI for 10min, and counted using a hemacytometer (Hausser Scientific). Finally, the remaining nuclei were spun at 4°C at 4000 rpm for 5 min, the supernatant was removed, and the pellet was flash frozen in liquid nitrogen. The nuclei were stored at −80°C until sonication.

#### IGN for Native ChIP-seq and ATAC-seq

Approximately 1.2 million worms were used per isolation for Native ChIP-seq, and 400,000 worms were used per isolation for ATAC-seq. After the final wash with M9, worms were washed with 10mL NPB. The supernatant was removed and the worms were resuspended in 6mL of M9 and transferred to a pre-chilled 7mL glass Dounce homogenizer. All of the Dounce and following steps were performed as for ChIP-seq except for the number of strainers. Worm debris was strained with three 30μm strainers and six 20 μm strainers. The samples were resuspended in 1mL NPB and left on ice until starting the desired genomic assay.

### ChIP-seq

For each sample, ∼20 million IGN were used. ChIP was performed as described previously (HAN *et al*. 2019). For each IP, 5μg of H3K27me3 antibody (Active Motif cat. no. 61018), 5μg of H3K36me3 antibody (Active Motif cat. no. 61022), or 5μg of anti-H3K9me3 (Abcam, #8898). Each IP was spiked with 90ng of *Drosophila* spike-in chromatin (Active Motif cat. no. 53083) and 0.5 μg of *Drosophila*-specific H2Av antibody was added to each sample.

For Native ChIP-seq, ∼12 million nuclei were used per sample. We adapted the protocol as described in (BRIND’AMOUR *et al*. 2015). Briefly, nuclei were washed in 1x PBS + 1x Complete Protease Inhibitor Cocktail EDTA-free and pelleted at 2000g for 10min. Nuclei were resuspended with 300 μL of Mnase buffer (50 mM HEPES pH 7.5, 110 mM NaCl, 40 mM KCl, 2 mM MgCl2, 1 mM CaCl2). Then 50 μL were aliquoted into 1.5 ml LoBind tubes(Eppendorf cat. no. 022431021). For each aliquot, 50 μL of Lysis Buffer 1 (1 mM Tris–HCl pH7.5, 1 mM PMSF, 1 mM DTT, 0.1% Tween 20, 0.2% NP-40, 0.01% digitonin) was added to each tube and incubated for 10 minutes. After the 10-minute incubation, 10U of Mnase (Sigma cat no. 10107921001) was added to each aliquot. Then 100μL of Mnase buffer was added, mixed by pipetting, and placed at 37°C for 4 min and 30 sec. Mnase digestion was stopped by adding 4 μL of 0.5M EDTA and then spun for 17000g for 10 min at 4°C. The supernatant was transferred to a fresh 1.5 mL LoBind tube. The pellet was resuspended in 250 μl of Lysis Buffer 2 (1 mM Tris–HCl pH7.5,1 mM PMSF, 1 mM DTT, 0.1% Tween 20, 1% Triton X-100 + 1x Complete Protease Inhibitor Cocktail) and then incubated for 2 hours while rotating at 4°C. Chromatin was pre-cleared with 40μL of Dynabeads protein G (Invitrogen cat no. 10003D) for 1 hour with rotation at 4°C. For input, 10% of lysate was set aside. For IP set up, 5μg of H3 antibody (Abcam cat. no. ab1791) was added. Each IP was spiked with 10ng of *Drosophila* spike-in chromatin (Active Motif cat. no. 53083) and 0.5 μg of *Drosophila*-specific H2Av antibody was added to each sample and incubated overnight with rotation at 4°C. The following day, 40μL of pre-blocked beads were added and incubated with rotation at 4°C for 2 hours. For the high salt wash, the ChIP samples with beads were washed 2 times with FA-150mM NaCl (50mM HEPES pH 7.5, 1mM EDTA pH 8.0, 1% Triton X-100, 0.1% Na deoxycholate, 150mM NaCl and 1x cOmplete proteinase inhibitor cocktail EDTA-free (Roche)) and 2 times with FA-500mM NaCl (50mM HEPES pH 7.5, 1mM EDTA pH 8.0, 1% Triton X-100, 0.1% Na deoxycholate, 500mM NaCl and 1x cOmplete proteinase inhibitor cocktail EDTA-free (Roche)). For the low salt wash, the ChIP samples were washed 2 times with FA-75mM NaCl (50mM HEPES pH 7.5, 1mM EDTA pH 8.0, 1% Triton X-100, 0.1% Na deoxycholate, 75mM NaCl and 1x cOmplete proteinase inhibitor cocktail EDTA-free (Roche)) and 2 times with FA-150mM NaCl (50mM HEPES pH 7.5, 1mM EDTA pH 8.0, 1% Triton X-100, 0.1% Na deoxycholate, 150mM NaCl and 1x cOmplete proteinase inhibitor cocktail EDTA-free (Roche)). The protein-DNA complex was eluted with 150μL of elution buffer (100mM NaHCO3, 0.2% SDS, 5 mM DTT, 1x TE) and incubated in the thermomixer at 65C for 15 min at 800 rpm. The elution step was repeated once more, and the supernatants were combined. For input, samples were brought up to 300μl with 1x TE. For both input and ChIP samples, 10μL of RNAse A (10mg/mL) was added and incubated in a thermomixer at 37°C for 15 min at 800rpm. Input and ChIP samples were digested with 3 μl proteinase K (20 mg/ml) at 55°C overnight.

For sequencing preparation, the input and ChIP samples were purified with Zymo ChIP DNA Clean & Concentrator kit (Zymo Research #D5205) according to the manufacturer’s instructions. The Yale Center for Genome Analysis (YCGA) prepared the library and performed sequencing. DNA integrity and fragment size were confirmed on a Bioanalyzer. Samples were sequenced on an Illumina NovaSeq using 100bp paired-end sequencing.

### ATAC-seq

For each sample, approximately 50,000 nuclei were used. The ATAC library was prepared as previously described (CORCES *et al*. 2017). Nuclei were washed with ATAC-Resuspension Lysis Buffer (10 mM Tris-HCl pH 7.4, 10 mM NaCl, 3 mM MgCl_2_, 0.1% Tween 20, and 0.01% Digitonin) then spun at 2000rpm for 10 min. Nuclei were then washed with ATAC-Resuspension Wash Buffer (10 mM Tris-HCl pH 7.4, 10 mM NaCl, 3 mM MgCl_2_, and 0.1% Tween 20). The nuclei pellet was resuspended in 50μl transposition mixture (2X Tagment DNA Buffer, 1X PBS, 0.01 % Digitonin, 0.1% Tween-20, 100 mM Tn5 Transposase) and incubated at 37°C for 30 min, shaking at 1000 rpm. The reaction was purified with Zymo DNA Clean and Concentrator-5 Kit (cat# D4014) according to the manufacturer’s instructions. DNA was pre-amplified with 2× NEBNext Master Mix (NEB M0541S). Library amplification was assessed by qPCR on Applied Biosystems QuantStudio 3 real-time PCR system. The library was purified with Zymo ChIP DNA Clean & Concentrator kit (Zymo Research #D4014) according to the manufacturer’s instructions. The Yale Center for Genome Analysis (YCGA) prepared the library and performed sequencing. DNA integrity and fragment size were confirmed on a Bioanalyzer. Samples were sequenced on an Illumina NovaSeq using 100bp paired-end sequencing.

### Computational analysis of genomic data

#### ChIP-seq processing

The quality of raw ChIP-seq data was assessed using FastQC (ANDREWS 2010), then mapped to ce11 with bowtie2 with default paired-end parameters (Langmead & Salzberg, 2012). To identify and filter for spike-in reads, raw ChIP-seq data were mapped to dm6 with bowtie2 with default settings. Whole animal histone ChIP-seq data were analyzed from available data: GSM3141776 and GSM3141788 (JÄNES *et al*. 2018), and GSM2515718 (MCMURCHY *et al*. 2017). These data were first mapped with bowtie2. The aligned reads were further processed by samtools to retain high quality reads (Q ≥ 30), and duplicate reads were removed by Picard MarkDuplicates. To minimize replication bias, bam files were downsampled to the size of the replicate with smaller library size, and then merged by samtools. For whole animal datasets, bam files were converted to bigwig tracks using deeptools bamCoverage -bs 50 –normalizeUsing CPM (RAMÍREZ *et al*. 2016). For spike-in normalization, the merged bam files were processed in R using ChIPSeqSpike package (DESCOSTES 2019). For Native ChIP-seq, the merged bam files were converted to bigwig tracks using deeptools bamCompare using input as control with parameters –normalize CPM -bs 50. Bigwig files were converted to z-score using custom R scripts. Peaks were called with MACS2 (ZHANG *et al*. 2008) with the parameters −q 0.05 –extsize 200. Metagene plots and heatmaps were generated using deepTools.

#### ATAC-seq processing

The quality of ATAC datasets were assessed with FastQC. Adapters were trimmed with cutadapt (MARCEL 2011) with the following parameters -a CTGTCTCTTATACACATCT. Files were aligned to ce11 with bowtie2 with the follow settings -X 2000 –no-mixed –no-discordant –very-sensitive. Bam files were processed for mapped quality Q ≥ 10, mitochondria reads were removed with samtools and duplicates were removed with Picard MarkDuplicates. Replicates were downsampled to minimize replication bias and then merged. Reads were then filtered for fragment size less than 100bp. Reads mapped to the + strand were offset by +4 bp and reads mapped to the – strand were offset by -5 bp. Bam files were converted to bigwigs by normalizing to RPM and converted to z-score using custom R scripts. Peaks were called using Genrich (https://github.com/jsh58/Genrich) with the following parameters -j -y -v. HOMER findMotifsGenome.pl with the parameter size -given was used to determine motif enrichment from ATAC peaks. The HOMER function annotatePeaks.pl was used to determine genes associated with differential peak analysis. Metagene and heatmaps were generated using deepTools.

#### Nucleosome occupancy

Merged bam files were used as inputs for NucleoATAC (SCHEP *et al*. 2015) using default settings. The occ.bedgraph files were used to visualize nucleosome occupancy. The NucleoATACR package was used to retrieve nucleosome occupancy, inter-dyad distance, and fuzziness scores, which were further analyzed using custom R scripts. Statistical significance was determined by Wilcoxon test.

#### Differential peak analysis

To identify differential peaks in the ATAC dataset, the Bioconductor package DiffBind (R. and D. 2011) was used with default settings. Filtered bam files and Genrich generated consensus peaks were used as inputs. Volcano plot was generated using custom R scripts.

#### piRNA expression analysis

piRNA expression data was obtained by publicly available data (GU *et al*. 2012; KASPER *et al*. 2014; BELTRAN *et al*. 2021). Reads were trimmed for adapters and aligned to ce11 with bowtie2. Reads matching piRNA annotations were matched and normalized with DESeq2. Custom R scripts were used to generate decile rank of piRNA abundance.

## DATA AVAILABILITY

The datasets supporting the conclusions of this article are available in Gene Expression Omnibus database under accession GSE232143.

## ACKNOWLEDGEMENTS

We would like to thank James Noonan and Bluma Lesch for equipment use, and members of the Reinke lab for valuable discussion and technical support. We thank Severin Uebbing for guidance on performing ATAC-seq and Matthew Simon and Bluma Lesch on discussion of ATAC analysis and Native H3 ChIP-seq. Sequencing service was conducted at the Yale Center for Genomic Analysis.

## AUTHOR CONTRIBUTIONS

V.R. and N.S. conceived the project. N.S. performed the experiments. N. S., L. E. G., and V. R. wrote the paper and discussed experimental results. N. S. performed deep sequencing data analysis. All authors read and approved the manuscript.

---

**Supplemental Figure 1.**
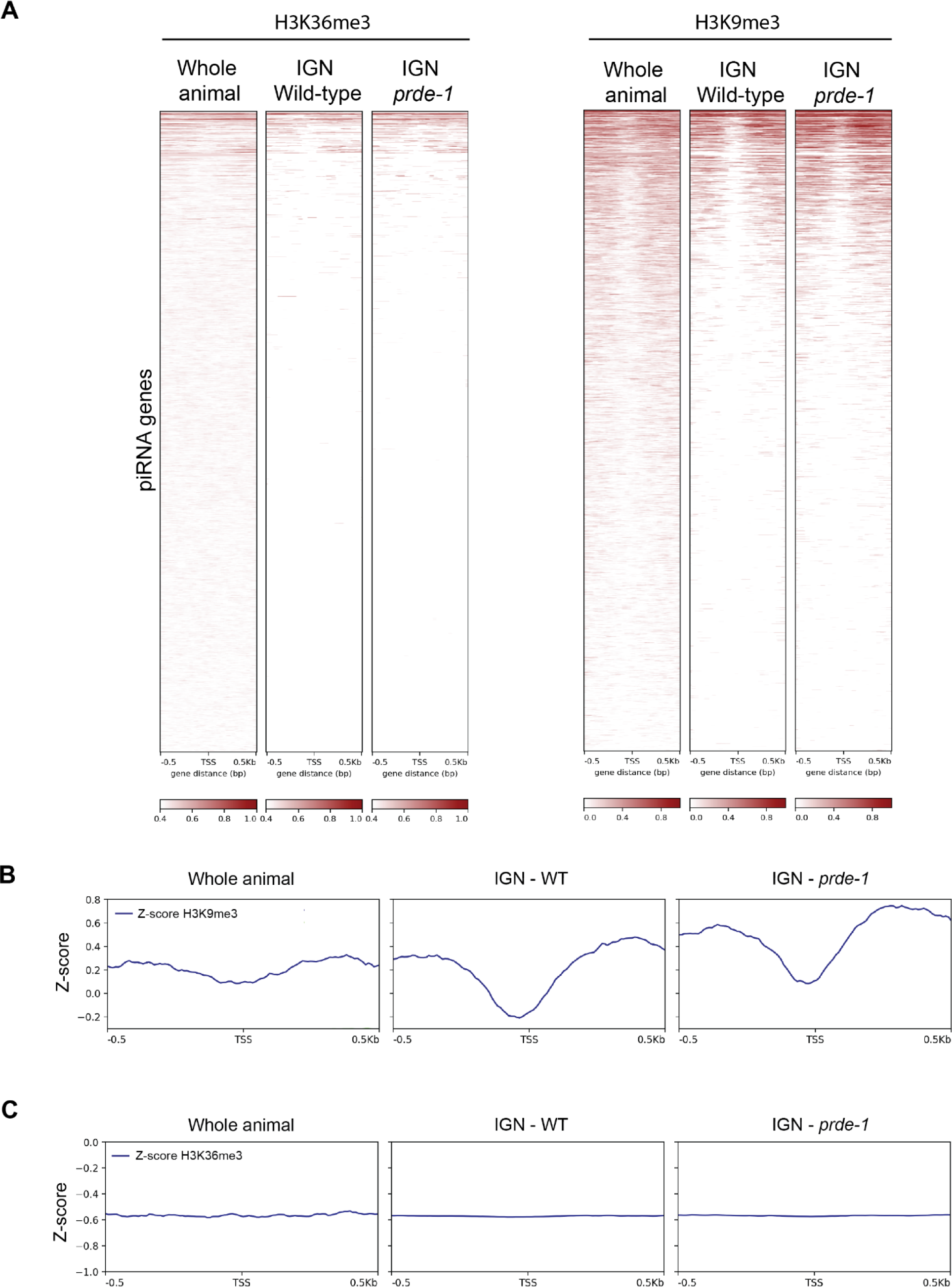
piRNA genes show low enrichment of H3K36me3 and H3K9me3 within germ nuclei. A) Heatmap of H3K36me3 and H3K9me3 levels centered on the transcription start site (TSS) of piRNA genes in piRNA clusters. The signal is represented as Z-scores. B) Metagene profile of H3K9me3 Z-score values across piRNA genes (1kb, centered on the piRNA TSS) located within H3K9me3 peaks. C) Metagene profile of H3K36me3 Z-score values across piRNA genes (1kb, centered on the piRNA TSS).

**Supplemental Figure 2.**
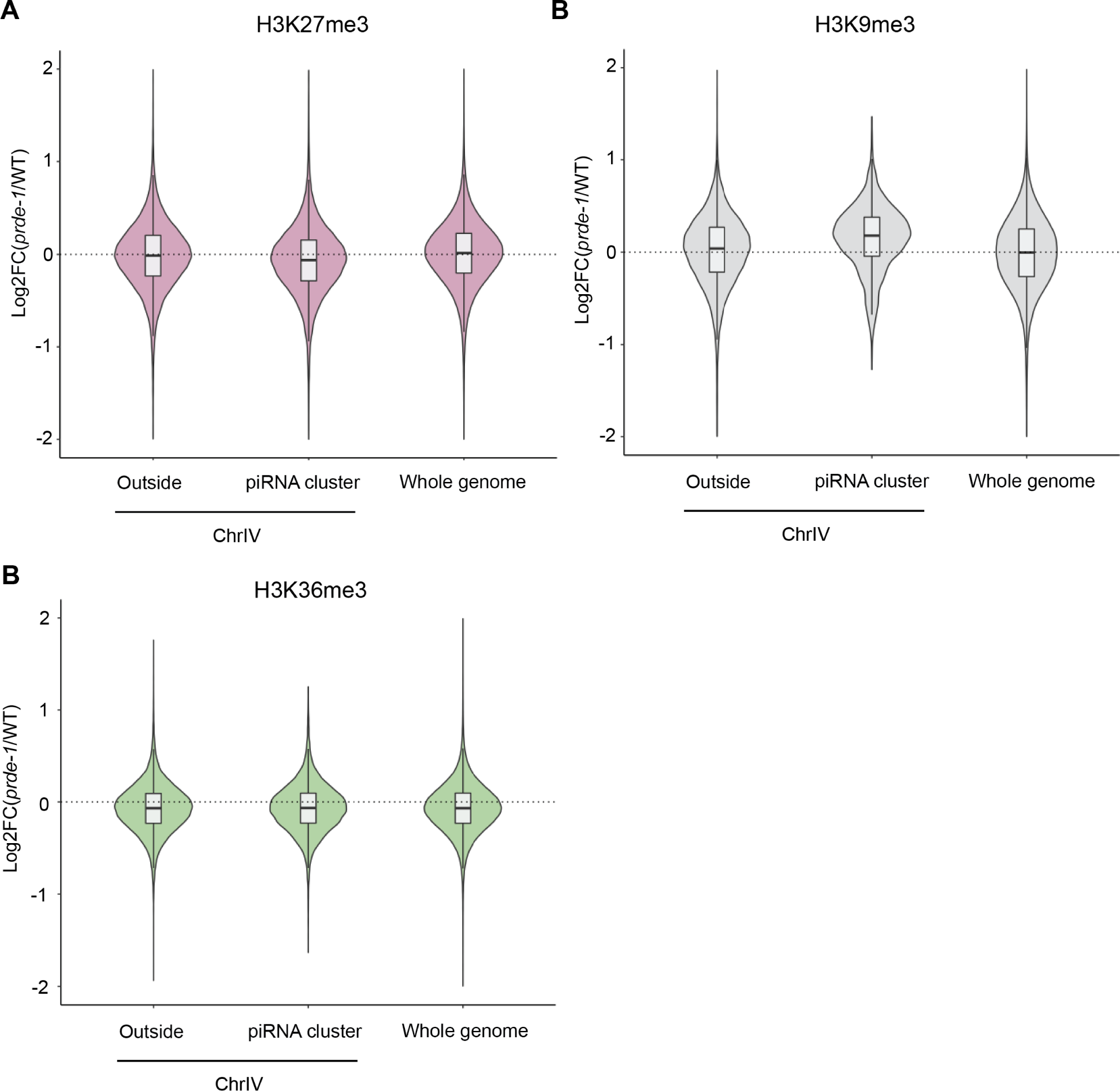
Subtle changes in repressive histone signals at piRNA clusters. Histone modification violin plots are: A) H3K27me3 B) H3K9me3 C) H3K36me3. Violin plots showing log2fold change in *prde-1* mutant relative to wild type of histone modification signal at their respective peaks. “Outside” refers to genomic regions on ChrIV that exclude piRNA gene cluster regions. “Whole genome” refers to genomic locations from all five autosomes and the X chromosome.

**Supplemental Figure 3.**
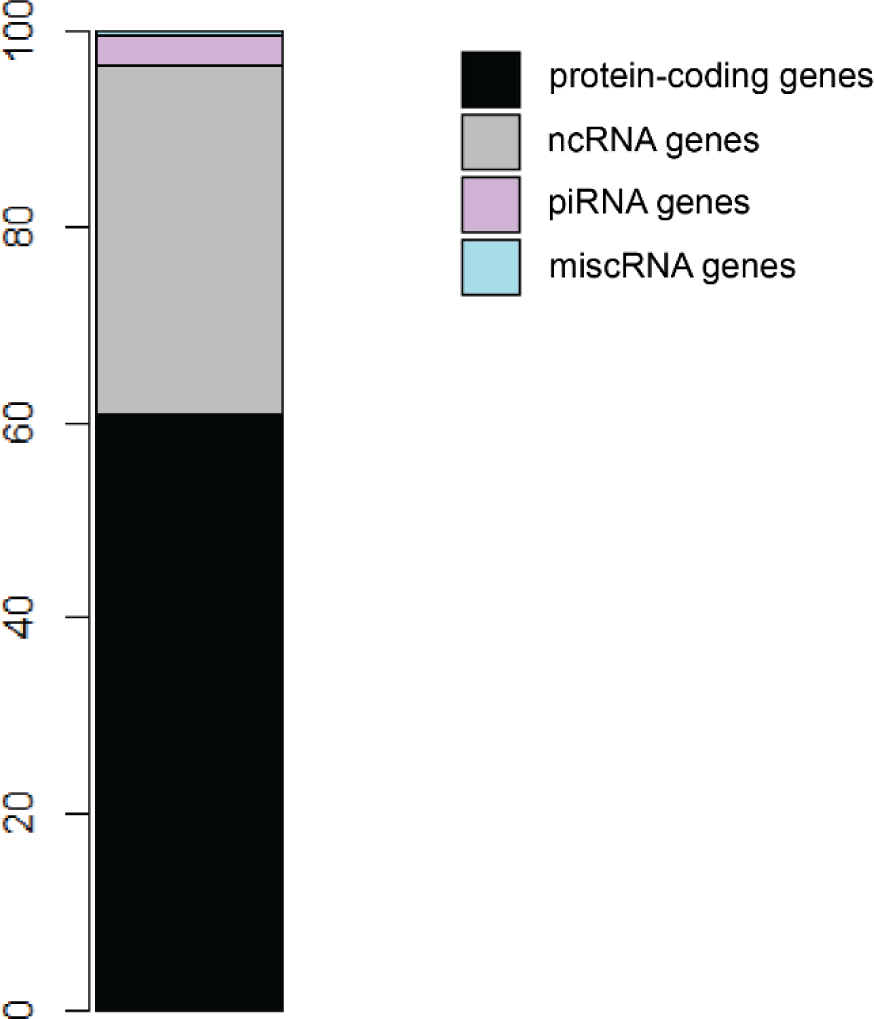
Differential chromatin accessibility peak loss is associated with protein-coding genes. The y-axis of the stacked barplot represents percentage of associated genes from peaks significantly lost in the *prde-1* mutant. Black represents protein-coding genes, grey represents noncoding RNA genes that exclude piRNA genes. Purple represents piRNA genes. Blue represent miscRNA genes.

**Supplemental Figure 4.**
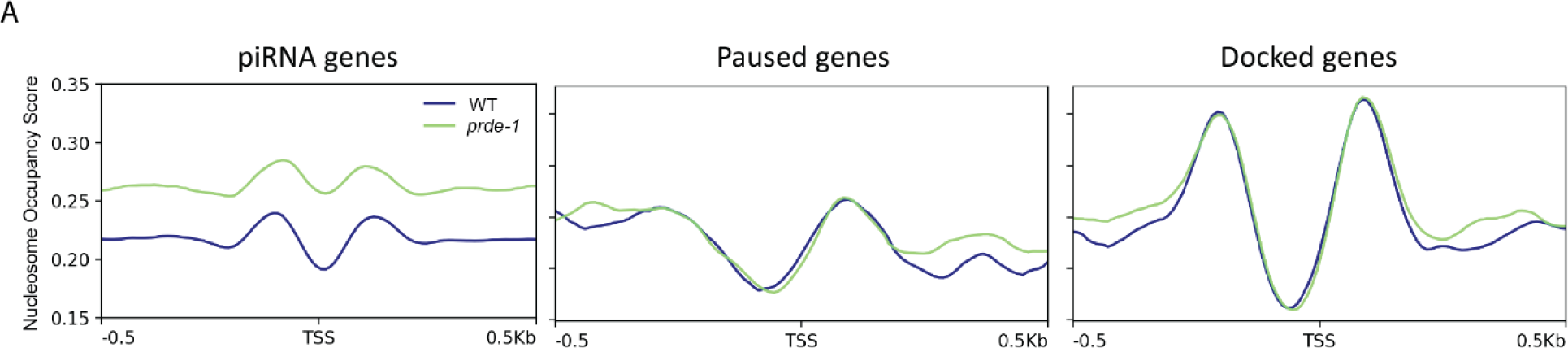
Loss of PRDE-1 causes no significant effect on known RNA Pol II paused and docked genes. A) Metagene analysis of nucleosome occupancy scores (1kb, centered on the TSS of piRNA genes). Wild type values are in blue while *prde-1* mutant values are in green. Middle and right plots represent nucleosome occupancy scores for known *C. elegans* RNA Pol II paused genes and genes regulated by RNA Pol II docked genes (Maxwell *et al* 2014).

